# A genetic perspective on Longobard-Era migrations

**DOI:** 10.1101/367201

**Authors:** Stefania Vai, Andrea Brunelli, Alessandra Modi, Francesca Tassi, Chiara Vergata, Elena Pilli, Martina Lari, Roberta Rosa Susca, Caterina Giostra, Luisella Pejrani Baricco, Elena Bedini, István Koncz, Tivadar Vida, Balázs Gusztáv Mende, Daniel Winger, Zuzana Loskotová, Krishna Veeramah, Patrick Geary, Guido Barbujani, David Caramelli, Silvia Ghirotto

**Affiliations:** Dipartimento di Biologia, Università di Firenze, 50122 Florence, Italy; Dipartimento di Scienze della Vita e Biotecnologie, Università di Ferrara, 44121 Ferrara, Italy; Fondazione Edmund Mach, 38010 San Michele all’Adige, Italy; Dipartimento di Storia, Archeologia e Storia dell’arte, Università Cattolica del Sacro Cuore, 20123 Milano, Italy; Soprintendenza Archeologia del Piemonte; Institute of Archaeological Sciences, Eötvös Loránd University, Budapest, Hungary; Research Centre for the Humanities, Hungarian Academy of Sciences, Hungary; Heinrich Schliemann Institut für Altertumswissenschaften Universität Rostock; Institute of Archaeology of the Czech Academy of Sciences, Brno; Department of Ecology and Evolution, Stony Brook University, 11790 Stony Brook, NY, United States of America; School of Historical Studies, Institute for Advanced Study, Princeton, New Jersey 08540, United States of America

**Keywords:** Population genomics, Ancient DNA, Migration, Approximate Bayesian Computation, Longobard

## Abstract

From the first century AD, Europe has been interested by population movements, commonly known as Barbarian migrations. Among these processes, the one involving the Longobard culture interested a vast region, but its dynamics and demographic impact remains largely unknown. Here we report 87 new complete mitochondrial sequences coming from nine early-medieval cemeteries located along the area interested by the Longobard migration (Czech Republic, Hungary and Italy). From the same locations, we sampled necropolises characterized by cultural markers associated with the Longobard culture (LC) and coeval burials where no such markers were found (NLC). Population genetics analysis and ABC modeling highlighted a similarity between LC individuals, as reflected by a certain degree of genetic continuity between these groups, that reached 70% among Hungary and Italy. Models postulating a contact between LC and NLC communities received also high support, indicating a complex dynamics of admixture in medieval Europe.

## Introduction

According to historical records, around the first century CE a Germanic population called “Longobard” was settled in the northern Elbe basin^1^. Around 500 CE the term “Longobard” reoccurs in the region north of the middle Danube, including Pannonia, and three generations later, in 568 CE, a people referred to as Longobards invaded and conquered much of Italy ^2^ The geographical spread of the word “Longobard” might evoke a quite large migratory phenomenon; however, the effective impact of the Longobard migration is still highly debated. There is a clear connection between the material culture of Pannonia and Italy at the end of the 6th century, suggesting strong interaction and communication – possibly as a result of a migration recorded in written sources – between these two regions (see eg.^3,4^). Archaeological and written sources, however, are open to different interpretations, and are unable to tell us whether such similarities in the material culture result from commercial exchanges or from migration. If the latter is the case, a further question arises, namely the relative contribution of immigrants and previously-settled people in the composition of the hybrid population. In particular, it is still highly debated whether and to what extent attributes such as grave goods and burial traditions are indicators of Lombard social identity, and whether the spread of these material markers across Europe is actually linked to population movements rather than to horizontal cultural transmission or trade. In this light, the analysis of ancient genetic data is fundamental to obtain a better understanding of past population dynamics and interactions.

The only ancient genetic data ever published from cemeteries associated with Longobard culture so far were sequences of the mtDNA control region from Piedmont, Italy^5,6^, and sequences of mtDNA control region and of informative positions in the coding region from the cemetery of Szólád in Hungary^7^. A broader analysis, based on larger assemblages of samples and genetic markers, is clearly necessary to provide a finer resolution of medieval populations movements and interactions in Europe. In this study we sequenced complete mitochondrial genomes from nine early-medieval cemeteries located in the Czech Republic, Hungary and Italy, for a total of 87 individuals. In some of these cemeteries, a portion of the individuals are buried with cultural markers in these areas traditionally associated with the Longobard culture (hereby we refer to these cemeteries as LC), as opposed to burial communities in which no artifacts or rituals associated by archaeologists to Longobard culture have been found in any graves. These necropolises, hereby referred as NLC, may represent local communities or other Barbaric groups previously migrated to this region. This extended sampling strategy provides an excellent condition to investigate the degree of genetic affinity between coeval LC and NLC burials, and to shed light on early-medieval dynamics in Europe.

## Materials and Methods

### Sample preparation and aDNA extraction

We collected teeth, postcranial elements and petrous bones for 135 individuals from nine early-medieval cemeteries located in the Czech Republic, Hungary and Italy (Fig. 1A, Table 1 and Table S1). Archeologically, 5 of these cemeteries have been partly associated with the Longobard culture: Mušov in the Czech Republic (LRCMUS), Szólád (LHSZ) and Hegykő (LHHEG) in Hungary, Collegno (LICOL) and Fara Olivana (LIFAR) in Italy. The other 4 sites -Fonyód (NLHFON), Hàcs-Béndekpuszta (NLHHACS) and Balantoszemes Szemesi-berek (NLHBAL) in Hungary, Torino-Giardini Reali (NLIGR) in Italy-while just preceding and being geographically close to the other cemeteries, did not show evidence of cultural practices traditionally termed Longobard. Additional informations on the archaeological context is available in the supplementary materials text. Bone samples were analyzed in the Molecular Anthropology Laboratory of the University of Florence, exclusively dedicated to ancient DNA analysis. Negative controls were used in all the experimental steps to monitor the absence of contaminants in reagent and environment. Long bones and tooth samples were cleaned by removing the surface layer using a dentist drill with disposable tips and exposed under UV light (λ = 254 nm) for 45 minutes on each side. 100 mg of powder were sampled from inside the compact portion of the bones and from the dentine of the tooth root and used in DNA extraction. Petrous bones were cleaned by brushing the surface and exposed under UV light (λ = 254 nm) for 45 minutes on each side. A disk saw was used to section the petrous bone and the inner surfaces were exposed under UV light (λ = 254 nm) before collecting the bone powder from the densest part of inner ear using a dentist drill with disposable tips as suggested in^8^. DNA was extracted following the protocol proposed in ^9^

**Table 1:**
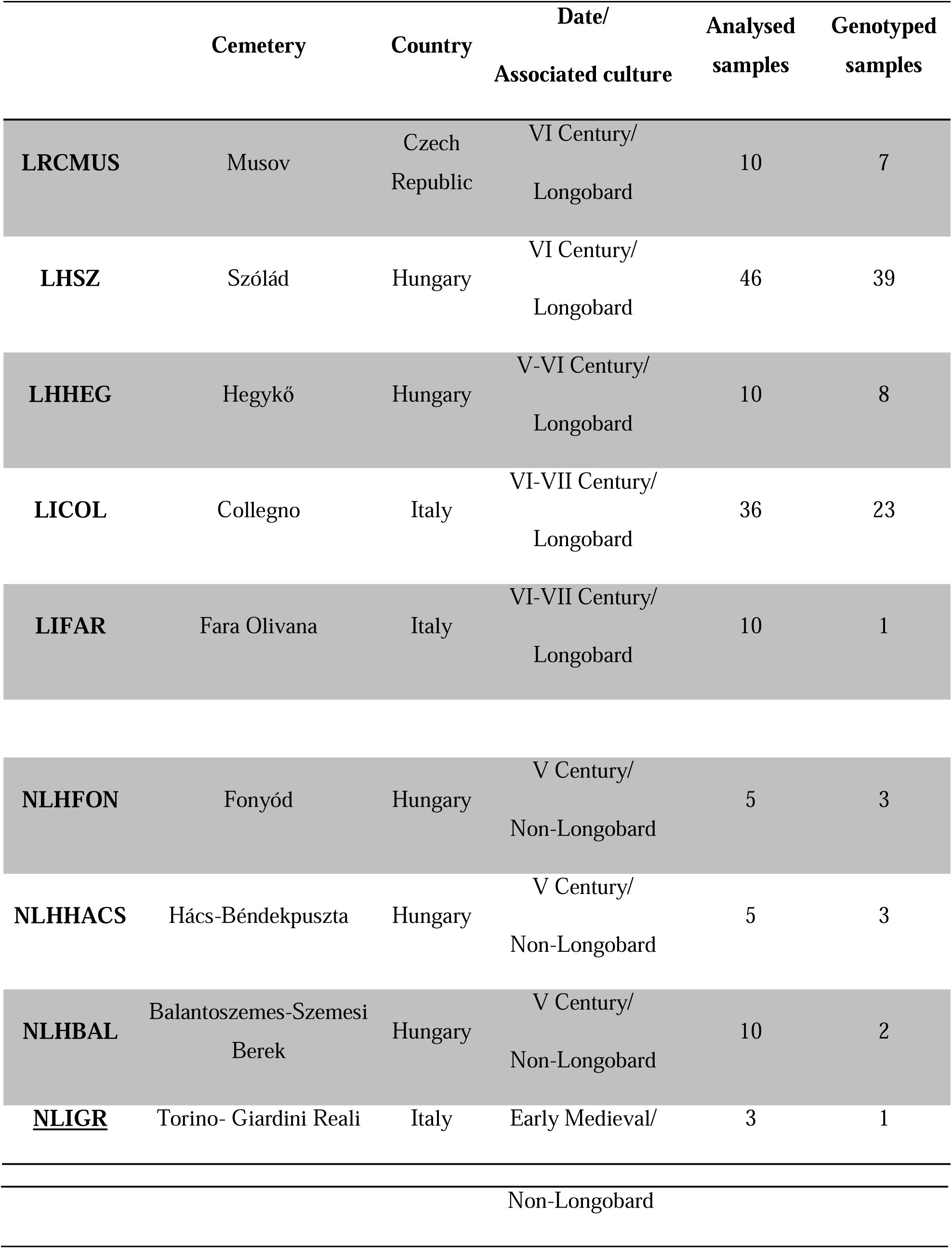
Cemetery information. For each group, geographical and chronological distribution and culture are reported. The table also shows the number of analyzed samples for each cemetery and the number of mitochondrial genomes obtained.

**Figure 1.**
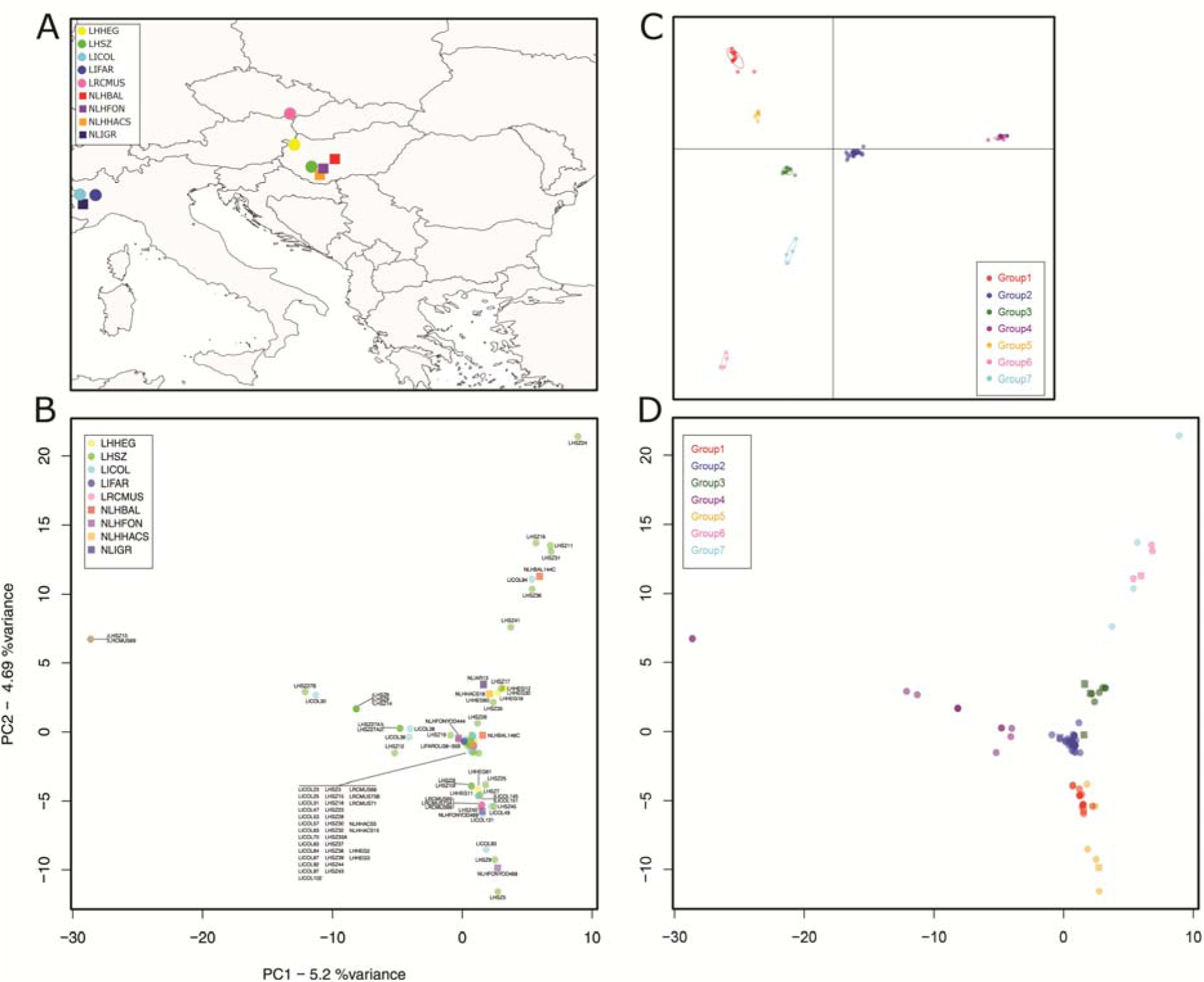
Geographical and genetic relationship between the newly sequenced individuals. (A) Location of the sampled necropolises. Here and through the other figures LC cemeteries are represented by a circle while NLC ones are indicated by a square. (B) PCA of the 87 newly sequenced LC and NLC individuals C) DAPC Scatterplot of the most supported K (7) highlighted by the *kmeans* analysis D) PCA where every individual is colored according to the groups located by the DAPC.

### NGS library preparation and sequencing

NGS libraries were prepared starting from 20 μl of DNA extract for each specimen following a double-stranded DNA protocol ^10^using a unique combination of two indexes per specimen. Libraries were enriched for mitochondrial DNA following a multiplexed capture protocol ^11^ and sequenced on an Illumina MiSeq run for 2×75+8+8 cycles.

### Mitochondrial DNA sequence pre-processing and mapping

Sequences were demultiplexed and sorted according to the sample, then raw sequence data were analyzed using the pipeline described in ^12^ Merged reads were mapped on the revised Cambridge Reference Sequence, rCRS (GenBank Accession Number NC_012920). Reads with mapping quality below 30 were discarded. Consensus sequence for each sample was obtained considering positions covered at least 3 fold, and base calling was performed with at least 70% of concordance between reads. Misincorporation pattern was analysed using MapDamage 2.0 ^13^ and contamination estimate was performed by contamMix ^14^ Only samples with at least 92% of the mitochondrial genome covered at least 3 fold, with CtoT values higher than 17% and MAP authentic values higher than 92% according to contamMix were considered for population genetics analysis. This resulted in a total of 78 suitable sequences from cemeteries archeologically associated with the Longobard culture (7 from LRCMUS, 39 from LHSZ, 8 from LHHEG, 23 from LICOL, 1 from LIFAR) and 9 suitable sequences from cemeteries not associated to the Lombard culture (3 from NLHFON, 3 from NLHHACS, 2 from NLHBAL and 1 from NLIGR) (table S2).

### Mitochondrial haplogroup definition and population genetics analyses

Mitochondrial haplogroups were assigned to the new mtDNA sequences according to PhyloTree Build 16 on Haplogrep ^15,16^. Phylogenetic networks were constructed using the Median Joining algorithm ^17^ implemented in Network 5.0 program (http://www.fluxus-technology.com). The 8 value was set to 0 and the transversions were weighted 3x the weight of transitions. Networks were subjected to maximum parsimony post-analysis. Haplogroup frequencies for medieval populations were retrieved from previously published data as outlined in ^18^. Since the vast majority of studies provided haplogroup frequencies inferred from HVRI, we reassigned the newly reported LC and NLC to haplogroups employing only this region. PCA based on haplogroup frequencies was conducted employing the function fviz_pca_biplot from the library factoextra ^19^ in R 3.4.0 ^20^. We compared LC and NLC complete mitochondrial genomes with a PCA using the function dudi.pca from the package adegenet ^21^. We also computed pairwise differences between sequences with Arlequin v. 3.5 ^22^ and visualized them employing an MDS computed with the function cmdscale. To locate possible population structure inside our dataset we assessed the best number of clusters inside it using the find.clusters function in adegenet ^23^, comparing the output of 10 independent runs using a custom-made R script. We then applied a DAPC analysis ^24^ on the dataset with 100,000 iterations.

### Demographic simulations

To explicitly test the importance of NLC populations on the genetic makeup of migrating LC individuals we conducted demographic simulations under an Approximate Bayesian Computation model framework. Accounting for the results of the exploratory analyses we first hypothesized two models called admixture and continuity (Fig. 3A/B). The former recreate a scenario where a migrant population, LC from northern Europe, receive gene flow from local NLC individuals before moving to colonize other regions. The continuity model instead postulates no contact between LC and NLC populations, mimicking only the proposed Longobard migration from Czech Republic to Italy. As Hungarian NLC populations contained individuals found buried along artifacts possibly associated with other Germanic cultures, we also developed two additional models named recent origin and recent origin+admixture. The recent origin model postulated a common ancestry between Hungarian NLC populations and LC ones sometimes around the 1st century CE), with a subsequent migration of the latter in Hungary. The recent origin+admixture model presents, intuitively, the same general structure depicted in the recent origin scenario with the exception of an event of admixture in Hungary between LC populations and NLC ones. In order to provide an unbiased representation of genetic diversity inside our dataset, we first removed related samples based on the kinship analysis presented in ^25^. We also excluded from our demographic dataset 2 individuals from Szólád (LHSZ27A1/2) as the dating of their burial suggest a more recent origin with respect to surrounding graves. This process resulted in a reduced dataset of 78 unrelated sequences. We obtained 20,000 simulations for each of the proposed scenarios with the software package ABCtoolbox ^26^. We performed the model selection procedure making use of the novel approach developed by Pudlo etl al. 2015, called ABC-rf^27^ which rely on the “random-forest” machine learning approach ^28^. Random forest uses the simulated datasets for each model in a reference table to predict the best suited model at each possible value of a set of covariates. After selecting it, another Random Forest obtained from regressing the probability of error of the same covariates determine the posterior probability. This procedure allows to overcame the difficulties traditionally associated with the choice of summary statistics, while gaining a larger discriminative power among the competing models ^28^. We built the reference table using the function abcrf from the package abcrf and employing a forest of 500 trees, as this number was suggested to provide the best trade-off between computational efficiency and statistical precision ^28^. We carried out the actual model comparisons and obtained the posterior probabilities of the winning models using the function predict from the same package. To summarize the genetic information contained in our sequences we considered the number of haplotypes, the number of private polymorphic sites, Tajima’s D, the mean number of pairwise differences for each population, the mean number of pairwise differences between populations and pairwise Fst. These summary statistics were obtained with arlsumstat ^22^ We validated the model selection procedure calculating the classification error through the abcrf function of the abcrf R package. To do this, we used as pseudo-observed datasets each dataset of our reference table (table S4). To verify whether the selected models were able to generate the observed data, we performed a linear discriminant analysis (LDA) using the function plot.abcrf. In order to estimate the parameters for the model chosen by the ABC-rf procedure (the admixture one) we ran further simulations reaching 1 million datasets. We applied a locally weighted multivariate regression ^29^ after a logtan transformation ^30^ of the 3,000 best-fitting simulations to estimate the admixture model’s parameter using an R scripts from http://code.google.com/p/popabc/source/browse/#svn%2Ftrunk%2Fscripts, modified by SG. The complete list of models’ parameters and of the associated prior distributions are presented in table S6.

## Results

### Medieval Mitochondrial genomes from Czech Republic, Hungary and Italy

The 87 medieval mitogenomes were sequenced to an average coverage depth of 86.10x (from 6.66x to 201.89x, Table S2). Sequences were assigned to 71 distinct haplotypes, all falling within the expected overall mitochondrial diversity of western Eurasian mtDNA (Table S2). Indeed, most individuals belong to the H, T2 and J lineages (respectively occurring in 33, 11 and 7 samples, Fig. 2B), all of them commonly observed in Europe ^31^. Mušov and Szólád LC cemeteries show similar frequencies of the H haplogroup (28% and 32% respectively), while in Collegno this frequency is doubled (60%) (Fig. 2B). The haplogroup distribution was rather different in Szólád and Hegykő (LC groups in Hungary); as an instance, we found the U8 haplogroup uniquely present in Hegykő, with a frequency of 25%. Phylogenetic links between haplotypes and their distribution among the archeological sites are shown in the Median Joining Network (Fig. 2A). Haplotypes were broadly grouped into their respective lineages and no general structure associated with geography or culture seems to be present.

**Figure 2.**
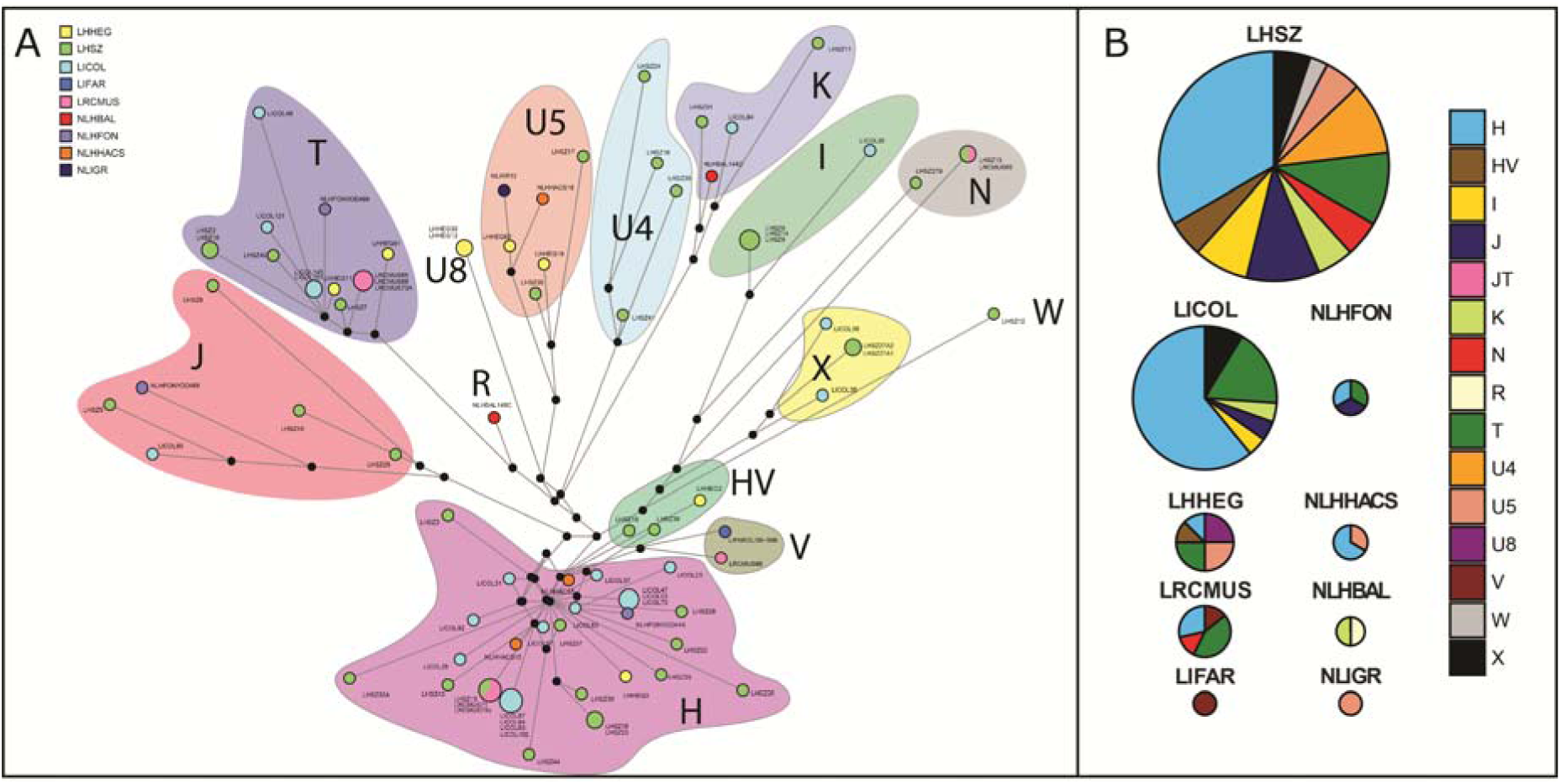
Haplogroup characterization of LC and NLC populations. (A) Median-joining network: the size of the circles is proportional to the number of samples carrying a specific haplotype, and the background shading indicates the affiliation of the lineages to the major haplogroups. (B) Pie charts representing the frequencies of the major haplogroups in the populations. The size of the chart is proportional to the number of samples inside each necropolis.

### Relationship between LC/NLC groups and other medieval samples

To elucidate the affinities between our new LC/NLC samples and coeval individuals we retrieved data on mtDNA haplogroup frequencies for 10 European medieval populations. A principal component analysis (PCA) on this expanded dataset highlighted the similarity between the LC graves of Szólád and medieval populations from Central Europe (Slovakia 800–1100 CE, Poland 1000–1400 CE) (Fig. S1). The Collegno individuals clustered midway between Slovakian and medieval samples from Southern Europe (Spain 500–600 CE, Italy 900–1400 CE).

### LC/NLC mitochondrial genetic variation and archaeological context

We explored the relationships between LC and NLC individuals through a principal component analysis (PCA, Fig. 1B). The first two axes of the PCA suggest a degree of similarity between groups, as NLC individuals are found across all of the range of genetic variation shown by the LC samples. There is also no clear geographical structure between samples in our dataset, with individuals from Italy, Hungary and Czech Republic clustering together. However, the first PC clearly separates a group of 12 LC individuals found at Szólád, Collegno and Mušov from a group composed by both LC and NLC individuals. The same pattern is also found when pairwise differences among individuals are plotted by multidimensional scaling (MDS, Fig. S2). To further investigate this peculiar genetic structure, we performed a *k-means* analysis and a discriminant analysis of principal components (DAPC) on the whole dataset. At K = 4 the 12 LC individuals located by the first PC form a cluster together, and stay together even at K = 7, the most supported number of clusters (Fig. 1C and 1D, Fig. S3). The presence in this group of LC sequences belonging to macrohaplogroups I and W, commonly found at high frequencies in northern Europe (e.g. Finland ^32^), suggests (although certainly does not prove) the existence of a possible link between these 12 LC individuals and northern Europe. The peculiarity of this group is strengthened by archaeological information from the Szólád cemetery, where 8 of the 12 individuals in this group originated, indicating that all these samples were found buried with typical Longobard artifacts and grave assemblages. We do not find the same tight association for the 3 samples from Collegno, where the 3 graves are indeed devoid of evident Germanic cultural markers; however they are not placed in a separate and marginal location—as for the tombs without grave goods found in Szólád —but among graves with wooden chambers and weapons. It is worth noting that weapon burials were quite scarce in 5th century Pannonia and 6th century Italy (e.g. Goths never buried weapons), and an increase in weapon burials started in Italy only after the Longobard migration. In this light, the individuals buried in this manner may have been members of the same community as well, but belonging to the lowest social level. This social condition could explain the absence of artifacts and could be related to mixed marriages, whose offspring had an inferior social rank. Finally, this group also includes an individual from the Musov graveyard. This finding is particularly interesting in light of the fact that the Musov necropolis has been only tentatively associated with Longobard occupation (see Supplementary Text for details), based on the presence of but a few archaeological markers.

### Testing models of Longobard migrations

To shed light on these early Medieval populations dynamics, we used an Approximate Bayesian Computation approach. We compared the possibility of admixture between LC and local NLC populations with a scenario postulating a simpler migration of LC individuals with no additional contact (Fig. 3A and 3B). We also tested two additional scenarios to explicitly consider that the similarities between LC and NLC individuals in Hungary may derive only from a recent common origin rather than local admixture (Fig. 3C and 3D). The admixture model received strong support, with probability of 86% (Table S3). The four models were well recognized by the ABC procedure we followed, as indicated by both the classification error (Table S4) and by the LDA plot (Fig. S4 and S5). This result can be interpreted as reflecting gene-flow from NLC inhabitants of the region into a migrating LC group. Indeed, when we estimated the extent of these admixture events, we observed that more than 80% of the genetic makeup of the Hungarian LC population could be traced to NLC people already inhabiting the region, while the Czech LC contributed around 18% (Table S5 and Fig. S6). This could either indicate a reduced contribution of Czech LC to the genetic make-up of the Hungarian LC, or that the Musov cemetery, while showing archaeological signs of Longobard occupation, is not a good proxy of the LC population that moved from the Czech Republic to Hungary. The Collegno individuals can, instead, trace more than 70% of their genetic makeup to LC populations migrating from Hungary, confirming the high degree of similarity shown by the exploratory analyses and supporting the migration hypothesis based on archaeological data.

**Figure 3.**
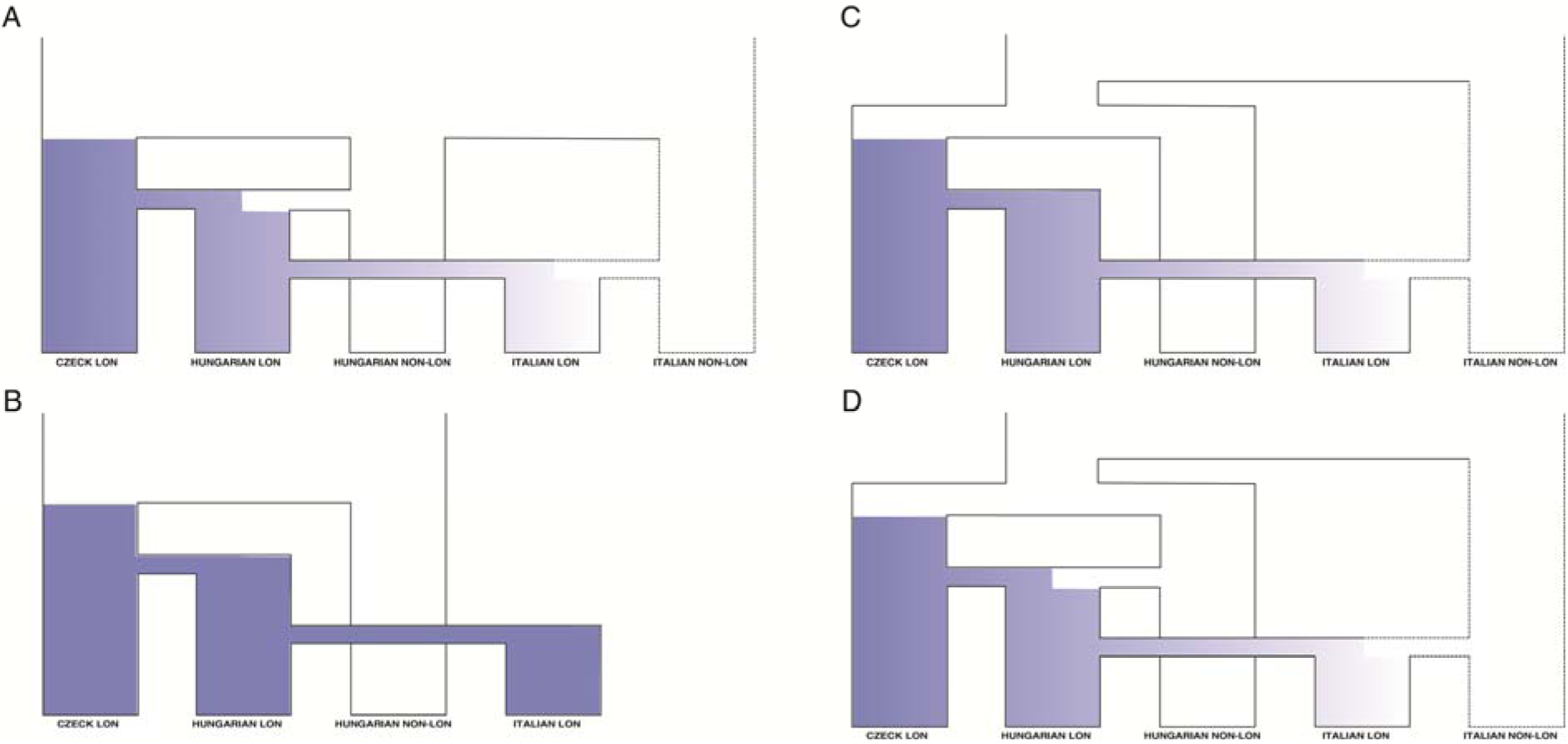
Alternative models of relationship between LC and NLC populations. A) admixture, (B) continuity, (C) recent origin, (D) recent origin + admixture. Continuous lines represent populations that were included in the simulations while dotted lines characterize ghost populations.

## Discussion

In this work, we extracted and analyzed complete mitochondrial genomes from 87 Early Medieval individuals sampled in 9 necropolises. Based on archaeological information we have classified these cemeteries as putatively occupied by Longobards or by different early Medieval communities. These genetic data have been used to explore, for the first time, the genetic variation and structure of these groups, so as to understand whether, and to what extent, different communities found along the route of the Longobard migration may resemble each other biologically.

Our explorative analyses highlighted a degree of genetic similarity between the LC and NLC communities, which was expected, given the well-known overall genetic similarity among all European populations. However, a peculiar set of samples, including only LC individuals from different European regions, appeared to be well distinct from the rest of the samples, consistent with the results of migrational exchanges between Pannonia and Italy. In most cases, these individuals were also associated with burials with typical Germanic grave goods, a term that here defines an assemblage of objects including Longobard artifacts, but not only them. This particular association, together with the presence in this cluster of haplogroups that reach high frequency in Northern European populations, suggests a possible link between this core group of individuals and the proposed homeland of different ancient barbarian Germanic groups, which will be further investigated. Therefore, at present we do not have evidence that the populations associated with northern Europe were not already in the region decades prior to the time that the Longobards entered the area. However these groups were likely related with the Gothic confederation, hence having different culture than Longobards. As both archaeological and bioarchaeometric analysis of the Szólád LC cemetery suggested that it was in use only for a short period of time, we suggest a recent arrival of LC communities in Hungary ^7^ The most interesting cases emerge when genetics can help to provide a better understanding in regard to social articulation of the Longobard groups, when the archaeological data are open to different interpretations. This happened, for instance, in Collegno, where the graves of these LC individuals were devoid of Longobard cultural markers, but placed among other burials rich of Longobard material culture. In this case, this pattern may suggest the presence of individuals from the lowest social level but still members of the Longobard community, rather than local/non Germanic people.

To better understand whether the observed genetic similarities across Europe could indeed result from migration, we estimated the evolutionary/demographic model best accounting for the degree of genetic variation presented by the LC and NLC samples here analyzed. Above and beyond the strong support observed for a model assuming admixture between LC and NLC populations (with respect to the models postulating recent common origin of Hungarian samples), the most interesting result was the high degree of genetic resemblance between the LC cemeteries in Hungary and Collegno, in Italy. We hence estimated that about 70% of the lineages found in Collegno actually derived from the Hungarian LC groups, in agreement with previous archaeological and historical hypotheses. This supports the idea that the spread of Longobards into Italy actually involved movements of fairly large numbers of people, who gave a substantial contribution to the gene pool of the resulting populations. This is even more remarkable thinking that, in many studied cases, military invasions are movements of males, and hence do not have consequences at the mtDNA level. Here, instead, we have evidence of changes in the composition of the mtDNA pool of an Italian population, supporting the view that immigration from Central Europe involved females as well as males.

While nuclear data would help elucidate these past population dynamics, the resolution provided by the mitochondrial genomes presented in this study, combined with detailed archaeological information, has allowed us to explicitly test demographic hypotheses, thus significantly improving our knowledge about different aspects of the complex pattern of migrations involving Longobard populations.

## Acknowledgments

We would like to thank Angela Livia Casadei for her help in the analysis of genetic data. This study has been supported by the following institutions and foundations: Riksbankens Jubileumsfond, Alexander von Humboldt Stiftung, Bundesministerium für Bildung und Forschung, Gerard B. Lambert Foundation, Università degli Studi di Padova, Institute for Advanced Study. KV and PG were supported by NSF #1450606. G.B., A.B. and F.T. were supported by the ERC Advanced Grant Agreement No 295733, ‘LanGeLin’ project.

## Data and Software availability

All newly generated data have been deposited in GenBank: MG182446 – MG182533.

## Supplementary information

Supplementary information is available at the European Journal of Human Genetics’s website

